# Repeated tDCS at clinically-relevant field intensity can boost concurrent motor learning in rats

**DOI:** 10.1101/2025.01.15.633248

**Authors:** Forouzan Farahani, Mihály Vöröslakos, Andrew M. Birnbaum, Mohamad FallahRad, Preston T.J.A. Williams, John H. Martin, Lucas C. Parra

**Affiliations:** Biomedical Engineering Department, City College of New York.; Molecular, Cellular and Biomedical Science, CUNY School of Medicine.; Neuroscience Institute, NYU Grossman School of Medicine, New York University.

## Abstract

Electric fields used in clinical trials with transcranial direct current stimulation (tDCS) are small, with magnitudes that have yet to demonstrate measurable effects in preclinical animal models. We hypothesized that weak stimulation will nevertheless produce sizable effects, provided that it is applied concurrently with behavioral training, and repeated over multiple sessions. We tested this here in a rodent model of dexterous motor-skill learning. We developed a preparation that allows concurrent stimulation during the performance of a pellet-reaching task in freely behaving rats. The task was automated to minimize experimenter bias. We measured field magnitudes intracranially to calibrate the stimulation current. In this study, only male rats were used. Animals were trained for 20 min with concurrent epicranial tDCS over 10 daily sessions. Behavior was recorded with high-speed video to quantify reaching dynamics. We also measured motor-evoked potentials (MEPs) bilaterally with epidural microstimulation. The new electrode montage enabled stable stimulation over 10 sessions with a field intensity of 2V/m at the motor cortex. The number of successful reaches improved across days of training, and the rate of learning was higher in the anodal group as compared to sham-control animals (F(1)=7.12, p=0.008, N=24). MEPs were not systematically affected by tDCS. Posthoc analysis suggests that tDCS modulated motor learning only for right-pawed animals, improving success of reaching, but limiting stereotypy in these animals. Repeated and concurrent anodal tDCS can boost motor-skill learning at clinically-relevant field intensities. In this animal model the effect interacted with paw preference and was not associated with corticospinal excitability.

**Significance Statement:** The effects of tDCS have been explored in numerous human clinical trials, but the mechanisms of action of weak electric fields remain unclear. In vitro studies show that constant electric fields at 2.5 V/m can enhance the efficacy of synapses undergoing plasticity. This study demonstrates in a rodent model that tDCS of only 2 Vm when applied concurrently to behavioral training can improve motor skill learning, and reduce stereotypy of reaching behavior. These effects accumulated over 10 days of training. Motor evoked potentials (MEP), which are often used to demonstrate plastic effects in humans on a time scale of hours, were not measurably affected by tDCS on this longer time scale.

## Introduction

Transcranial Direct Current Stimulation (tDCS) utilizes low-intensity electrical current delivered to the brain through electrodes positioned on the scalp (Bikson et al., 2019). The effects of tDCS have been explored in numerous human clinical trials (Chase et al., 2020; Chen et al., 2022; Lefaucheur et al., 2017), but the mechanisms of action of such weak electric fields remains unclear.

The predominant theory posits that long-term effects of tDCS are mediated by effects on synaptic plasticity (Stagg & Nitsche, 2011). *In vitro* studies have demonstrated the effectiveness of anodal DCS in boosting synaptic plasticity (Farahani et al., 2021; Fritsch et al., 2010; Gellner et al., 2020; Kronberg et al., 2017; Ranieri et al., 2012; Rohan et al., 2015; Sun et al., 2016) by increasing neuronal firing (Farahani et al., 2021). However, the electric fields used in these investigations are relatively large, with fields of at least 5 V/m and up to 20 V/m. In contrast, in human tDCS experiments, typical scalp currents of 2mA electric fields can reach at most 1V/m (Huang et al., 2017; Opitz et al., 2016). Most *in vitro* studies have first applied stimulation in a single session and then tested its effect on synaptic plasticity thereafter (Barbati et al., 2020; Podda et al., 2016; Ranieri et al., 2012). Such “offline” effects may be less effective as compared to applying tDCS concurrently with synaptic plasticity induction (Kronberg et al., 2017, 2020; Sharma et al., 2022). Additionally, one may gain in effectiveness by leveraging spaced learning, whereby repeated bouts of training can boost learning (Smolen et al., 2016). We have found cumulative effects of DCS *in vitro* when paired with repeated synaptic plasticity induction (Sharma et al., 2022). Cumulative effects of tDCS are also leveraged in human experiments delivered in daily sessions over 5 to 15 days or more (Boggio et al., 2007; Ho et al., 2016, pp. 3-; Loo et al., 2012). We hypothesized that plasticity induced *in vivo* over multiple training sessions with concurrent stimulation should produce observable enhancements of behavioral and physiological markers of plasticity, despite weak electric fields.

The *in vivo* animal studies to-date leave significant ambiguity as to the actual electric field intensity achieved within the brain. While human and non-human primate experiments have measured and modeled intracranial electric fields (Huang et al., 2017; Opitz et al., 2016; Vöröslakos et al., 2018), similar calibration efforts have not yet been performed in rats. Despite numerous *in vivo* studies of tDCS in rodents (Barbati et al., 2020; Cambiaghi et al., 2010; Liu et al., 2019; Monai et al., 2016; Podda et al., 2016; Rohan et al., 2015), measurements of electric fields are rare (Yu et al., 2023), including in the context of motor skill learning, which is the focus of this study (Barbati et al., 2020; Longo et al., 2022; Ramanathan et al., 2018). Based on our measurements (Farahani et al., 2024) and other previous studies (Yu et al., 2023), we estimate that *in vivo* experiments in rodents have used fields of 15 V/m or more. These field intensities may surpass the threshold required for neuronal firing and one expects that effects are categorically different from what may be achieved with sub-threshold weak fields used in human tDCS studies. Therefore, there remains a significant gap between mechanistic *in vitro* and *in vivo* animal work at 5-20V/m and human experimentation at less than 1V/m. By directly measuring intracranial electric fields in the rodent brain, we aim to calibrate the electrical current, so as to approach clinically realistic electric field magnitudes.

In humans, tDCS has been explored in particular in the context of motor skill learning (Amadi et al., 2015; Fritsch et al., 2010; Hsu et al., 2023; Reis et al., 2009; Yamaguchi et al., 2020). Motor skills improve through repeated practice, and this improvement might be increased when learning is combined with tDCS. Structural and functional changes in the primary motor cortex (M1) have been linked to motor skill learning (Kleim et al., 1998; Rioult-Pedotti et al., 2000; Xu et al., 2009). We previously postulated that tDCS paired with training will boost the specific effects of training by enhancing Hebbian synaptic plasticity (Kronberg et al., 2020). To investigate the long-term effects of concurrent tDCS with training, we decided to focus this study on its impact on motor learning over repeated sessions. We chose the single-pellet reaching task, which is commonly used when studying motor skill learning in rodents over the course of 10 or more daily training sessions (Ellens et al., 2016; Ramanathan et al., 2018; Salameh et al., 2020). It involves training rodents to perform a sequence of movements that include reaching, grasping, and retrieving a single pellet. The task has similarities to the act of humans reaching for food (Sacrey et al., 2009).

The plastic effects of tDCS on motor cortex are often discussed in the context of motor evoked potentials (MEP) (Stagg & Nitsche, 2011), which measure corticospinal system excitability (Pellicciari et al., 2013). Anodal tDCS has been shown to increase MEPs in both human and animal studies (Cambiaghi et al., 2010; Nitsche et al., 2004; Nitsche & Paulus, 2000, 2001). These studies have investigated the impact of tDCS on MEP applied either while the subject is at rest (Nitsche & Paulus, 2000; Pellicciari et al., 2013) during task performance (Amadi et al., 2015; Ambrus et al., 2016; Antal et al., 2007; Wiltshire & Watkins, 2020) or before/after a task (Amadi et al., 2015; Yamaguchi et al., 2020). However, it is not clear whether changes in TMS evoked MEP are long-lasting and directly related to motor learning. Indeed, the literature on MEP and motor learning is mixed. There are human and animal studies showing that motor learning expands the cortical areas that can evoke an MEP (Brus-Ramer et al., 2009; Monfils et al., 2005; Pascual-Leone et al., 1995). However, studies linking motor skill learning to MEP amplitudes have shown mixed results (Ambrus et al., 2016; Wiltshire & Watkins, 2020).

The aim of our study is to bridge the gap between human studies and in vivo animal experiments by utilizing a low-intensity electric field, similar to that used in human studies. In rodents, we intend to measure MEPs as a possible marker of plastic effects. To achieve repeated tDCS concurrent with training, we have designed a novel electrode montage that ensures electrochemical stability and animal safety while maintaining stimulation intensity across 10 days of training. To ensure an intended 2V/m electric field in the M1 region of the brain, we intracranially measure field intensity and calibrate the electrical current. We use high-speed video and automated tracking software (Bova et al., 2019) to analyze the stereotypy of reaching behavior. This is the first *in vivo* animal study to intentionally limit the intensity of the electric field and to apply tDCS concurrently with behavioral training across several training sessions.

## Methods

### Animals

All animal procedures were approved by the ASRC Institutional Animal Care and Use Committee At The City College of New York, CUNY (protocol 2020–5). This study used 27 male Long-Evans rats (300-400 g) housed in pairs on a reversed light/dark schedule; 24 animals were used for the motor training protocol, and 3 for the measurement of electric field intensity. Food and water were available ad libitum except during reach training and testing, as described below.

### Chest and epicranial electrode implantation for tDCS

In this experiment, rats were instrumented with tDCS electrodes on the skull (epicranial) and chest (Fig. 2A). The chest electrode was made of a platinum grid and implanted (Fig. 2B, 10 mm x 10mm, Goodfellow 512-248-13). We tested several epicranial electrodes (Fig. 2B-D), but for the main experiment, we selected a platinum plate (3mm x 3mm) placed inside a gel-filled pocket permanently fixed on the epicranial (Fig. 2D).

For electrode implantation surgeries, the rat was anesthetized with isoflurane (3-4% induction, and 1.5-2% maintenance) in oxygen. Heart rate, respiration rate, and oxygen saturation were monitored throughout the surgery, and deep anesthesia was verified by non-response to tail pinch. The surgical sites were shaved, cleaned and infiltrated with a local anesthetic (bupivacaine injection max 2%). A sagittal midline incision (1 cm in length) of the scalp was made for the epicranial electrode, and a sternal sagittal incision (3 cm in length) was made for the chest electrode.

A cable was tunneled subcutaneously between the openings from the chest to the left occipital corner alongside the neck. The electrodes were attached at each end. The chest electrode grid was fixed to the pectoral fascia at the four opposing corners using non-absorbable suture and the edges were coated with silicone to avoid tissue abrasion. Finally, the skin was closed using cutaneous sutures.

To affix the epicranial electrode, the animal was then placed in a stereotactic frame. The epicranial periosteum was scraped off and the bone was cleaned with saline and 3% H2O2 (Fritsch et al., 2017). Permanent resin cement (Kerr) was then applied to the bottom rim of the electrode holder. The midpoint of the holder was positioned 3 mm lateral from the bregma contralateral to the preferred paw and +1.5 mm anterior from the bregma. After the holder was fixed in place, we ensured that the stimulation area was free of resin cement using a drill. The pocket holder was filled with conductive gel (Signa Gel-electrode gel), a 3×3 mm^2^ platinum plate (soldered to the uninsulated platinum wire) was placed in the pocket, and the lateral border was sealed with a small drop of cement. Contact resistance was measured to ensure it was below 10 kOhm, which would indicate that cement is occluding the contact area, but not in the range of a few Ohms which would indicate inadequate insulation of the pocket.

Electrode connectors (Plastic One E363/0) were embedded in a sealed headcap on the vertex of the skull formed with dental acrylic and anchored to the skull by setting four nylon screws. The skin incisions were sutured and treated with topical antibiotic ointment. Animals were given 7 days of recovery after surgery, and before commencing the motor training with concurrent tDCS.

### EMG electrode implantation and recording

After 10 days of training with concurrent tDCS, rats were evaluated for the strength of motor evoked potential (MEP) responses in a key forelimb muscle for reaching. We used an epidural electrode to stimulate a large portion of the forelimb motor cortex which activates short-latency muscle response. This required removing the epicranial electrode and performing a craniotomy over the motor cortex to place a bipolar stimulation electrode on the dural surface. Anesthesia was induced with ketamine (70 mg/kg i.p.) and xylazine (7 mg/kg i.p.) and the rat was placed in a stereotaxic frame. The skin was incised along the midline, and the head cap was removed. A craniotomy (4 mm by 2 mm) was made over the motor cortex of each hemisphere, spanning from bregma (AP 0) to bregma +4 mm and from 1 mm lateral to the midline to 3 mm lateral.

The stimulation electrode consisted of two parallel wires (2 mm length, 1mm spacing, Product info) placed on the exposed dura. We used an isolated stimulator (AM Systems Model 2100) to deliver pulses (0.2 ms pulses repeated 3 times at 300 Hz) through the bipolar epidural electrodes. The minimum current intensity (in the range of 0.5-4mA) to evoke a small MEP in 90% of trials (above the noise floor of 0.05mV) was set as the movement threshold. MEP recruitment was evaluated from 20 trials tested at 90%, 100%, 120%, 140%, and 200% of motor threshold intensities. The procedure was then repeated by placing the stimulation electrode on the other hemisphere.

This was repeated on the opposite hemisphere to record evoked EMG signals from both motor cortices. This terminal experiment lasted for approximately 3 hours, during which time the animals were monitored for the absence of reflexes every 15 minutes with a tail pinch, and Ketamine was re-administered if the animal reacted to the pinch.

Evoked potentials were recorded in the ECR muscles bilaterally, i.e., trained and untrained forelimb. Muscle activity was recorded differentially with pairs of microwire hook electrodes (PFA-coated tungsten wires, 500 µm., A-M Systems, Inc) inserted percutaneously in the extensor carpi radialis of the left and right forelimb. EMG placement was confirmed in the ECR muscle by applying biphasic pulses (0.05-0.2 mA) to evoke wrist dorsiflexion. The raw EMG signals were amplified (Model 1700), low-pass filtered at 5 kHz and sampled at 10 kHz with CED 1401.

### MEP analysis

We calculated the mean amplitude of the rectified MEP signal in a time window of 10 - 30 ms after the first pulse of stimulation and then considered the median across the 20 trials as MEP amplitude (Extended Data Fig. 4-1). Of the 24 animals that were trained, two animals were lost during the MEP procedure, and one animal was lost on day 8 of training as the headcap detached. Additionally, we excluded one animal due to unreliable MEP threshold values, and another due to dominant motor unit activity, leaving 19 animals for MEP analysis. We performed all analyses at the 140% threshold, and confirmed similar results at different epidural stimulation intensities. We used the logarithm of MEPs in our statistical analyses to approximate normal distributed values. Therefore, an analysis of differences in MEP amplitude constitutes an analysis of amplitude ratios between conditions.

### Application of epicranial tDCS

To achieve 2 V/m in the primary motor cortex, we administered an electric current with an intensity of 150 μA (calibrated based on electric field measurements, see below). We maintained a fixed duration of 20 minutes for each experiment in addition to a 10-second linear ramp-up and ramp-down. We used a current-controlled stimulator (Caputron LCI 1107 High Precision) to generate this tDCS waveform. Throughout the stimulation period, we used an ampere meter to monitor the current intensity. If the stimulator could not maintain the required current we considered the electrode contact lost and removed the animal from the experiment. This happened on day 8 for the first animal tested with the 2 mm platinum plate, and occurred only once in 24 animals of the main experiment using the 3 mm electrode, also on day 8. The stimulation was delivered to the rat via a wire connected to a double brush commutator (P1 Technology), ensuring the rat could move freely in the chamber. We performed the same surgery to implant electrodes for the animals in the control group; however, no current was applied during the training sessions in these control animals.

### Alternative tDCS instrumentation

We tested several electrode montages to address a number of limitations in previous in-vivo rodent experiments with tDCS: first, the stimulation montage should ensure the delivery of electrical current at a desired field intensity; second, the montage should provide stable current across multiple sessions; and third, it should allow rats to move freely in the chamber and be unencumbered during reaching. We tested and dismissed a “jacket” to hold the cathode (Oh et al., 2019), because the jacket restricted the rats’ mobility. Instead, we sutured the electrode on the chest (Fig 2B) (Fritsch et al., 2017), using a grid electrode for optimal electrochemical stability. Importantly, we observed no signs of discomfort in the animal’s behavior when the current was applied.

For the anode, we tested a conventional electrode holder glued to the skull (Fig 2C), which allows for the replacement of gel and Ag/AgCl electrodes (Ethridge et al., 2022). However, infections developed on the skull over 10 days of treatment. We also evaluated a permanent gel/electrode enclosure attached to the scalp (Fig 2D) -- the pocket described above, which was previously developed (Vöröslakos et al., 2018). Here we enlarged the size of the pocket and electrode to maintain electro/chemical stability for 10 training sessions of 20 minutes with a current of 150 μA.

### Electric field measurement

We measured the electric field generated by transcranial currents in 3 animals that did not participate in the motor training protocol. To record electric fields a 4-channel device was created as follows. Tungsten wires of 50 μm in diameter and 3 cm in length were prepared by removing the insulation from one end of each wire. Two such wires were inserted into a stainless-steel tube (26-gauge needle shortened to 3 mm), positioned 5 mm from the tube’s end and separated by 1 mm from each other, and glued in place. A metal screw was affixed to the skull at the occipital bone to serve as the ground electrode and covered with dental cement for mechanical support. This ground wire was connected to the same connector combining two 2-channel single shank devices creating a grounded single device with 4 contacts in a square planar configuration with a 1mm side distance. To connect the device to a preamplifier headstage, a header pin connector was soldered to an Omnetics adapter. Electric potentials measured at the 4 contacts were digitized at 20 kS/s using an RHD2000 recording system with a 32-channel preamplifier (C3314, Intan technologies, Los Angeles, CA).

For the experiment, a craniotomy was performed on the temporal bone at a depth of 3 mm from the skull’s top and 1.44 mm anterior to bregma, and the dura was removed (following similar procedures as above). The 4-channel tungsten device was inserted to a target depth of 2.4 mm from the lateral brain surface (Fig. 2E). Transcranial current was applied with varying frequencies (10, 100, and 1000 Hz) and intensities (10, 20, and 40 µA) through the skull/chest electrodes while electric potentials were measured in the 4 channels. The potential difference between the four channels divided by the 1mm distance captures the electric field magnitude and direction in 2D. Measuring gain as 0Hz is technically challenging because electrode impedance effects play an important role for constant currents. Fortunately, a previous study (Opitz et al., 2016) has shown that gains vary only slightly across frequency (10% drop from 1 Hz to 100 Hz). We selected higher frequencies because the recording equipment we used here is designed for recording high frequency spiking activity and is calibrated for unit gain at 1000 Hz.

### Behavioral chamber and training

We constructed a fully automated reaching chamber using polycarbonate panels (Fig 1A), following a previous publication (Bova et al., 2019). The front panel of the chamber has a small opening through which rats can access a sugar pellet (Fig 1B). An infrared sensor located at the back of the chamber detected the presence of the rat and triggered the movement of a pedestal to move a new pellet up into the reaching position. We used high-speed video recording at 310 fps (Basler ace acA2000-340kc) to capture frontal and side views with a single camera using three mirrors (Fig 1B). Another infrared sensor at the front detects the presence of the paw, and the camera records 200 frames before and 800 frames after the detection from a memory buffer (StreamPix). The pedestal was programmed to move out of position one second after the paw is detected at the front to end the trial.

**Fig 1:**
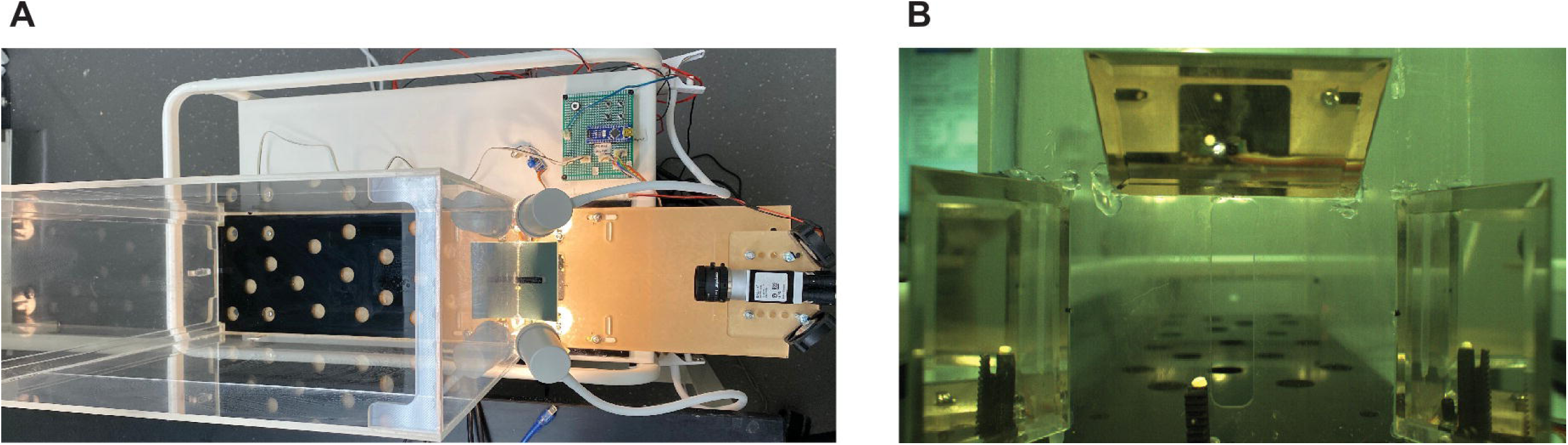
Training chamber and video recording. **A:** Overhead view of the chamber apparatus for automated reach training. Each trial required the rat to traverse to the rear end of the chamber to trigger a sensor (infrared beam) that initiates automated loading of a food pellet on a pedestal into target position outside an aperture The pedestal is offset to the side of the opening to be sure it can only be reached with the preferred paw. **B:** Arrangement of three mirrors to capture the movement of the rats’ paws in 3D. High-speed video recording is triggered by another sensor outside the opening.

The experimental timeline is shown in Fig. 3A and follows these steps in sequence:

**Fig 2:**
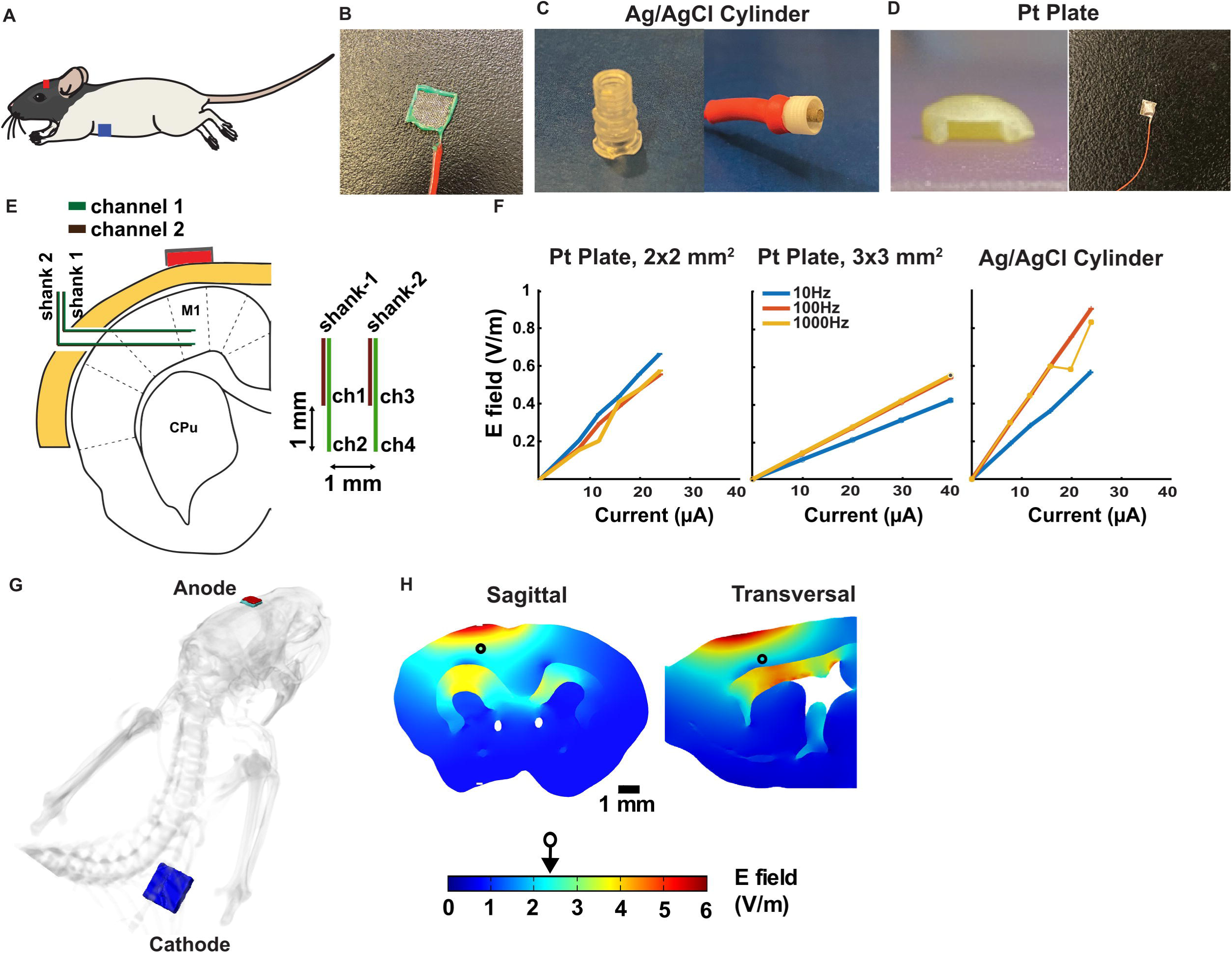
**A:** Schematic of the montage (Adapted from (Tang, 2020)) **B:** Cathode grid electrode implanted in the chest. **C:** Left: conventional epicranial electrode holder with a radius of 3 mm, right: Ag/AgCl stimulation electrode. **D:** Left: custom-made epicranial electrode holder made from dental cement, right: stimulation electrode 3×3 mm^2^ platinum plate. **E:** Recording shanks with two contacts per shank forming a rectangle of 1 x 1 mm to measure the E-field in 2D. **F:** Measurement of the electric field amplitude (combining vertical and horizontal components) at 10, 100, and 1000 Hz at different current intensities. These were measured with different electrode holders in different animals (one animal per stimulation configuration). **G:** Anatomically detailed current flow model of the current montage as described in (Farahani et al., 2024) **H:** Model of electric field magnitudes when stimulating with a current of 150µA applied continuously over the 20 min of stimulation. The increased intensity in deeper tissue is due to increased resistivity assigned to this white matter structure.

**Fig 3:**
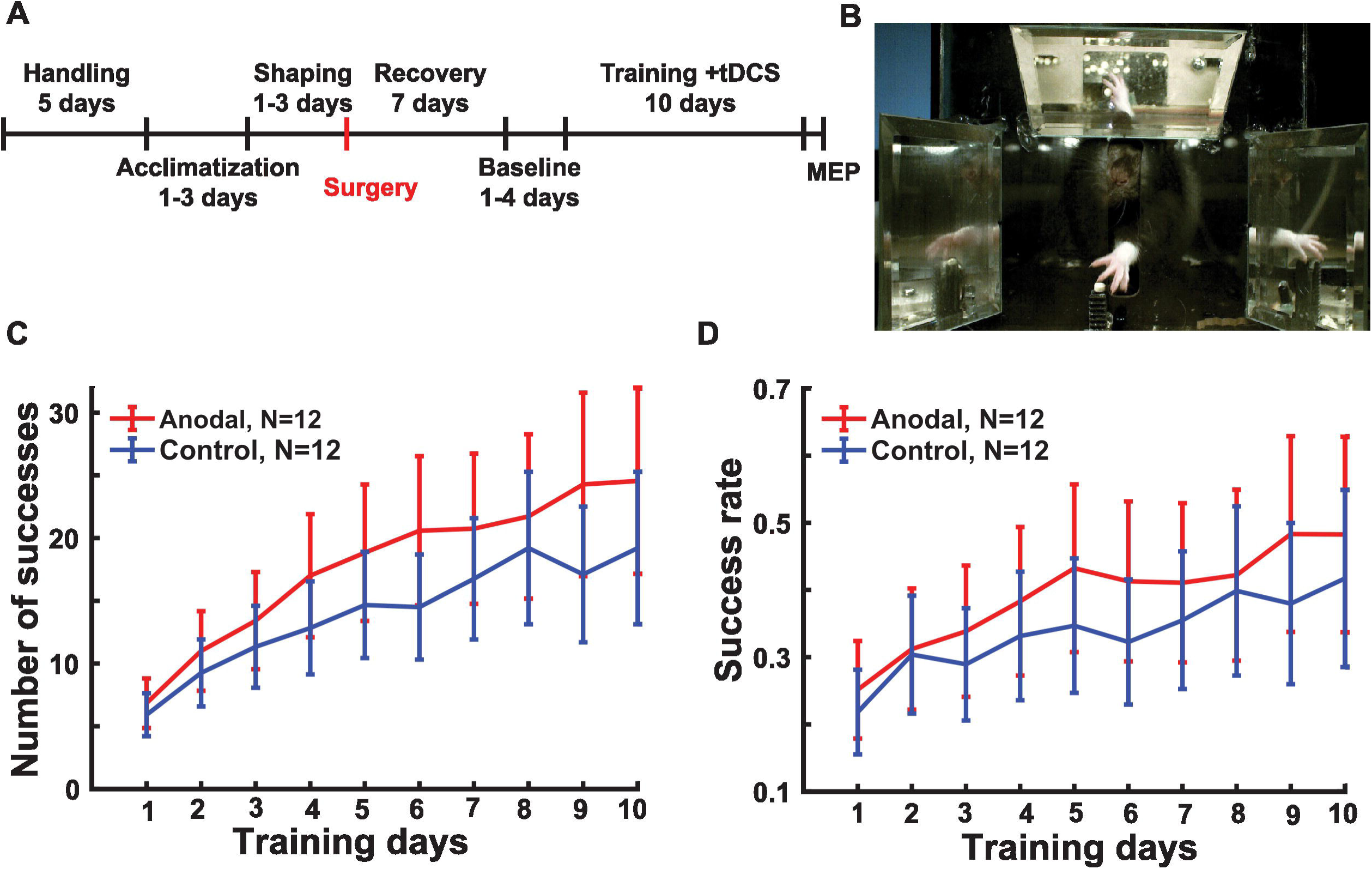
Effect of tDCS on motor skill learning. **A:** Experiment timeline from acclimatization, surgery, training and MEP recording. **B:** Frontal image of the rat reaching for a food pellet (white). Mirrors on top, left and right facilitate 3D recording of the paw motion with a single video camera. **C:** Number of successful reaching attempts across days of training. Lines indicate mean (and SEM) across animals in the anodal (red, N=12) and control groups (blue, N=12). **D:** Success rate across days of training.

#### Handling (5 days)

First, rats are handled for 15 minutes each day for at least five days to become accustomed to the experimenter.

#### Acclimatization (1-3 days)

The rats are introduced to the sugar pellets in their home cage. After putting the rats on food restriction (targeting 10-15% weight loss within 1 week), we placed sugar pellets in both the front and back of the reaching chamber and observed the rat’s behavior for 15 minutes. We use forceps to hold the sugar pellet through the opening on the front panel and allow the rat to eat the pellets from the forceps a few times.

#### Shaping (1-3 days)

As the rat attempts to eat the pellet, we gradually pull the forceps back until the rat learns to grasp the pellet with its paw. We repeat this step 10 times to determine the rat’s paw preference and align the pedestal accordingly. We consider the paw that a rat uses 6 out of 10 times as the preferred paw.

We placed a pellet on the pedestal and encouraged the rat to attempt grasping it with the baited forceps several times. Once the rat reaches for the pellet ten times without being baited, we move on to the next step. After placing the rat in the chamber, we bait it to the back of the chamber using a sugar pellet. As the rat reaches the back of the chamber, it breaks the infrared beam, and the pedestal is automatically loaded with a sugar pellet and raised into reaching position. We repeat this process several times until the rat learns to move to both the back and front of the chamber without being baited. This period of shaping is followed by surgery and recovery, where food restriction is paused.

Baseline (1-3 days): After recovering from surgery the animal is acquainted with the task under food restriction until it can reach for the pellet successfully at least once.

#### Training (10 days)

Automated training takes 10 days for 20 minutes and 20 seconds with concurrent tDCS. To evaluate the performance of each rat, we count the number of successful reaches in this constant time interval, i.e. the animal grasped the pellet and brought it to its mouth without dropping it.

### Video analysis and stereotypy of grasping movement

We recorded the movement of the paw with high-speed video while the rats reached for the food pellet. Video recording is triggered by a light sensor at the start of the reach trial lasting from 350 ms to 2.5 s. The videos have a frame rate of 309 fps with a resolution of 2032×1086 pixels. Following the recording, videos were compressed using ffmpeg (Tomar, 2006), on average, by a factor of ∼75. We used DeepLabCut (Mathis et al., 2018) to track the position of the wrist and digits in these compressed videos in 2D. The trajectory tracking model of DeepLabcut underwent training on 10 videos, featuring a single right-pawed rat, with one video from each session. In each video, 19 frames were manually labeled, marking six points of interest: five digits, wrist, and pellet (Fig. 5A). This training data was split into a 95% train set and 5% test set. Training for 1,030,000 iterations resulted in a 1.26-pixel training error and a 4.12-pixel testing error. For consistency, videos of left-pawed rats were horizontally flipped before analysis, enabling the utilization of the same model for all rats.

The model outputs horizontal and vertical positions for each of the 6 points (Fig. 5B). The resulting traces were median-filtered with a 15 ms (5 samples) window. Animals tend to grab several times within a single video recording (trial). To detect instances of an individual grab within a trial we used a template matching technique (Fig. 5-2). This entailed creating a template trajectory (Fig. 5-2B), cross-correlating it with the entire trajectory of the trial (Fig. 5-2C), and selecting time points with peak correlation corresponding to individual grabs (Fig. 5-2D). Templates of 150 ms duration (50 samples) were formed for each animal by averaging trajectories from three grabs in each training session. Cross-correlation was performed separately for each of the 10 coordinates and averaged across all 10 (vertical and horizontal coordinates for each of the 5 digits. The last peak in a trial at the end of the video was considered the final grab and the correlation value at that time point was taken as a measure of stereotypy. The various analytic choices to measure stereotypy were blinded for tDCS condition, paw preference, and number of successful trials.

Successful reaches involve not just the grab but require the animal to guide the sugar pellet successfully into their mouth. The number of successful trials was counted visually on the recorded videos and this counting was also blinded to the tDCS condition. The mirrors showing depth of reaching (Z direction) helped identify successful reaches during visual inspection. Code to load and visualize all the reaching trajectories can be found at: https://github.com/birnybaum/Rat-Reaching-Task.

### Statistical analysis

We used mixed effect models to analyze the number of correct grasps, MEP amplitudes and stereotypy. Days were coded on a logarithmic scale to account for saturating response. 3 animals were missing days 8-10, which is handled conventionally by the mixed effect model. Effects of tDCS that accumulate over days of training are reflected in an interaction between days and the stimulation condition.

## Results

### Stimulation configuration and calibration of the electric field intensity

In our hypothesis, the strength of the field intensity at the target is a crucial determinant of effectiveness. To measure this field intensity, we recorded voltage in M1 at four points in a square planar latice of 1 mm spacing (Fig 2E). We applied sinusoidal transcranial stimulation at 3 different frequencies (10, 100 and 1000 Hz) with different current intensities (Fig 2F). Except for some outlier measurements, the field increased linearly with current intensity, as expected. The gain (V/m per μA) is consistent for 100 Hz and 1000 Hz and somewhat lower at 10 Hz. This suggests that the non-uniform gain observed here is due to the recording equipment, which is calibrated for 1000 Hz. As expected, different epicranial electrode holders result in different gains (0.0214, 0.0141, and 0.0367 V/m/µA with the frequency of stimulation at 100 Hz). For calibration of the field, we used the gain observed at 1000 Hz with the 3×3 mm^2^ platinum electrode, which was 0.55 V/m/μA. Thus, a 150 μA results in an estimated electric field of 2 V/m, which is closer to the range of human studies. Computational modeling with this electrode montage (Fig. 2G) conducted in a parallel study (Farahani et al., 2024) shows approximately 2V/m at the depth corresponding to M1 (Fig. 2H).

### Anodal tDCS boosts the performance in motor skill learning

We predicted that the concurrent application of tDCS during training can enhance the learning performance. In other words, a higher slope of the learning curve for the animals which received anodal tDCS or interaction of tDCS with days of training (Fig 3C). A linear mixed effect model for the number of successful reaches finds an interaction between days and tDCS (t(227)=2.68, F(1)=7.12, p=0.008, N=24). There was also an obvious effect of days as the learning progressed (t(227)= 9.64, F(1)=267.64, p=1.2×10^-18^. There was no significant effect of tDCS (t(227)=0.23, F(1)=0.0530, p=0.81, N=24), indicating that the benefits of tDCS accumulate in the course of training with equal performance on day 1.

To determine if the cumulative effect is the result of increased accuracy or speed, i.e. a larger number of attempts in the fixed 20 min, we inspected the success rate per pellet presented (Fig. 3D). The success rate is the ratio of successful reaches to total reaches, expressed as a fraction. While it is numerically larger for the anodal group, the effect is not significant in this follow-up exploratory comparison (test same as above, interaction F(1,277)=2.82, p=0.094). The same is true for the number of reaching attempts (F(1,277)=0.26, p=0.61). This finding suggests that performance gains resulted from an increase in both speed and accuracy, which together resulted in a significant improvement of successful reaches.

### MEPs are not affected by training for tDCS

We aimed to determine whether there was a lasting impact on MEPs when anodal tDCS was applied concurrently with motor training. We hypothesized that training will increase corticospinal excitability and pairing anodal tDCS with training will further boost this increase. To test this, we measured MEP amplitudes (Extended Data Fig. 4-1) following 1-2 days after the final day of training. We measured in both trained and untrained paws with pulsed cortical stimulation of ipsi- and contralateral motor cortices (Fig. 4A; 4 measures in total). As is customary, we measured MEP at multiple epidural stimulation intensities (Fig. 4B).

**Fig 4:**
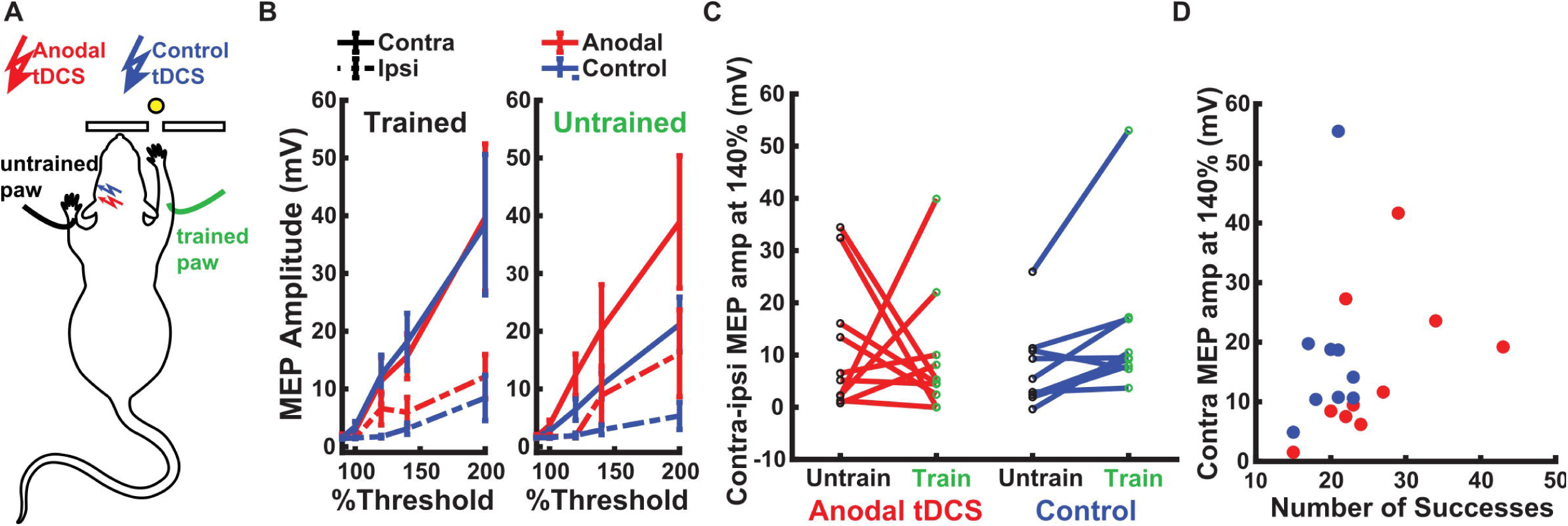
Motor Evoked Potentials (MEP) **A:** MEPs are measured with electrodes placed on the trained (green) and untrained wrist extensor muscles (black). The “trained” paw is the one with which the animal learned to grab the food pellet (yellow) through a narrow slit. tDCS electrode is placed over the contralateral motor cortex of the trained paw during 10 days of training. In terminal MEP experiments, bipolar electrodes are placed in direct epidural contact over both left and right motor cortices (ipsi- and contralateral sides relative to recording electrodes). (Extended Data Fig. 4-1 shows examples of MEP traces) **B:** MEP amplitude measured in trained and untrained paw with ipsi and contralateral epidural pulsed stimulation. Motor recruitment curves can be obtained by adjusting the pulse intensity as a percentage of motor thresholds. Lines indicate mean (and SEM) across animals in the anodal (red, N=10) and control groups (blue, N=9). **C:** MEP amplitude of trained and untrained paw in response to stimulation of the contralateral hemisphere at 140% of threshold. Each line is one animal. **D:** MEP amplitude at 140% threshold in the trained paw with contralateral epidural pulsed stimulation vs. behavioral performance at the end of learning (number of successful reaching attempts on the 10th day). Each point is one animal.

First, basic anatomy suggests that stimulating the contralateral hemisphere would result in a greater MEP amplitude compared to the ipsilateral side, regardless of training or tDCS. We used a linear mixed-effect model with a fixed-effect factor of side (contralateral vs ipsilateral epidural stimulation) and animal as a random effect. As expected, this analysis shows that contralateral MEP was larger than ipsilateral MEP (solid lines > dashed lines in Fig. 4B, t(74)= 9.17, p=7.7 x 10^-14^).

We predicted that MEP amplitudes will increase with training for the trained paw as compared to the untrained paw, and that this increase is larger in animals that received anodal tDCS, as compared to the control group. We next analyzed the log-difference in MEP between contra- and ipsilateral epidural stimulation.As both use the same recording electrode/paw that controls for variations in EMG strength across animals. We used a linear mixed-effect model with fixed-effect factors of tDCS (control vs anodal) and training (trained vs untrained paw), and animal as a random effect. Contrary to our prediction, we did not find an effect of training (t(34)=1.3, p=0.18) or tDCS (t(34)=-1.63, p=0.11) on MEP amplitude, and we did not observe an interaction between the two (t(34)=1.29 p=0.20) (Fig. 4C). However, analysis in the control animals alone, does reproduce the expected effect of training, i.e. the trained paw had stronger MEP than the untrained paw (t(16)=2.25, p=0.03). We also expected that MEP will correlate with performance across animals. Again, we used a mixed effect model for the log-amplitude of MEP, this time with fixed factors of success and tDCS (Fig. 4D). We did not see any effect of success (t(72)=0.86, p=0.39), nor tDCS (t(72)=-0.79, p=0.42) on MEP amplitude (Fig. 4D).

To rule out that the lack of an effect with tDCS is due to variability in threshold currents we tested whether threshold current affects MEP amplitude and found no effect (fixed effect of current intensity at 100%) with rats as a random effect: t(74)=-1.25, p=0.13). To summarize, while training seems to have enhanced MEPs, we did not see an effect of tDCS.

### Behavior becomes more stereotypical for right-pawed rats

Next, we conducted an exploratory analysis of behavior using high-speed video recordings. We expected the movements during reaching to become more stereotypical with training, and this effect to be enhanced in the anodal group. We used DeepLabCut to label and track the digits of the paw in the video during reaching in 2D (Fig. 5A). Reaching attempts often involved multiple grabs. We selected the last grab in each trial because it included all the successful reaches (example trajectory for one grab in Fig. 5B). Post hoc analysis using the first grab gave similar results, albeit less clear. Examples of the last grab from multiple trials are shown for one animal (Fig. 5C first session, 5D last session, the time course in X and Y direction; Extended Data Fig. 5-2). We measure stereotypy as the correlation coefficient of the reaching trajectories with that of a template (Fig. 5-2).

**Fig 5:**
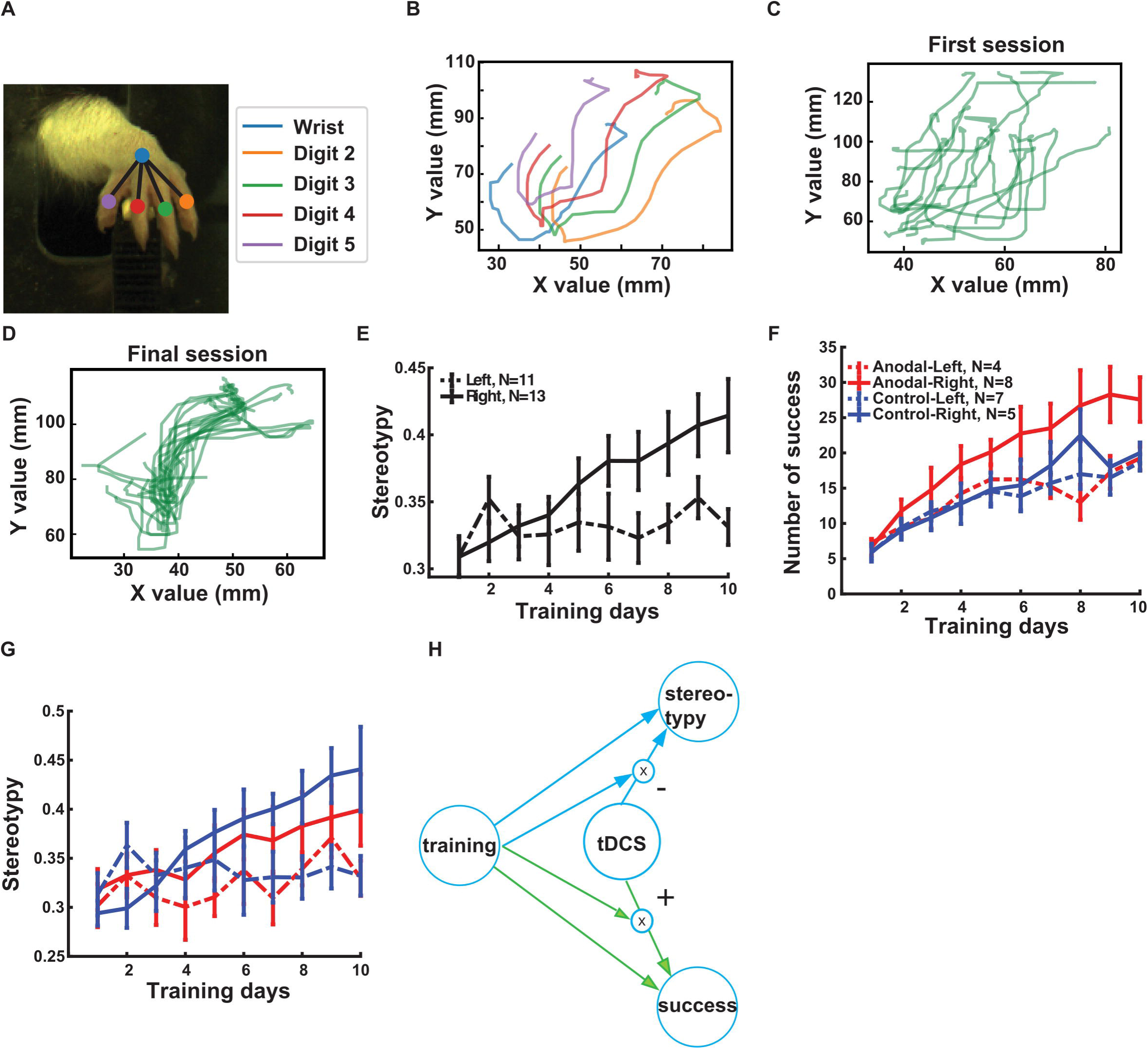
Effect of motor training and tDCS on stereotypy. **A:** A snapshot of a paw with the colored dots indicating the position labels. **B:** Sample trajectories for each digit in a single grab. **C:** Reaching trajectories of the middle digit (#3) across several trials in the first session for a right-pawed rat. **D**: Reaching trajectories of the middle finger across several trials in the last session for the same rat. **E:** Stereotypy across training sessions for right-pawed and left-pawed animals (solid and dashed lines respectively; error bars indicate SEM). (Extended Data Fig. 5-2 shows reaching trajectories for 4 digits as a sample) **F:** Number of successes separating now by paw preference and stimulation conditions (red anodal tDCS and blue control). **G:** The same as F but for stereotypy. **H:** Schematic of the results of statistical analysis for right-pawed animals. Green is the planned primary outcome measure and blue is the secondary measure. (Extended Data Fig. 5 −1 shows schematic of the results of statistical analysis including paw-preference as a factor)

We noted in the videos a marked difference in behavior between left and right-pawed animals. Whereas right-pawed animals mostly used a single paw for reaching, left-pawed animals occasionally also reached with the right paw, or used the right paw to support body position. Overall their behavior before and after the reaching movement was more diverse.

We therefore inspected stereotypy taking paw preference into account. Of the 24 animals, 13 had a right-paw preference, roughly in line with existing literature (Manns et al., 2021). In the present study, it appears that only right-pawed animals increased in stereotypy (Fig. 5E). It also appears that tDCS diminished the effect of training on right-pawed animals (Fig. 5F). To avoid cumbersome 3-way interactions, we first present here results focusing on animals with right-pawed preference, and show results including all animals in Extended Data Fig. 5-1. We used a mixed effect model to test for a fixed effect of training (day) and tDCS (anodal/control) as well as their interaction, and with animals as a random effect. We find in right-pawed animals that training improves stereotypy (t(120)=5.54, p=1.8×10^-7^), and that tDCS reduced this effect (interaction: t(120)=-4.09, p=7.6×10^-5^), but there was no effect of tDCS in isolation (t(120)= 1.12, p=0.26) (Fig. 5G). Separate analysis of successful and unsuccessful trials revealed that this result is primarily driven by the unsuccessful trials.

The results of stereotypy prompted us to perform the same exploratory analysis of paw preference also on reaching success. Fig. 5F suggests that only right-pawed animals benefited from tDCS. Again, we focused analysis on right-pawed animals, with results on all animals in Fig. 5-1. We tested for a fixed effect of training (day) and tDCS (anodal/control) as well as their interaction. Additionally, we included a fixed effect of stereotypy, as we expected that stereotypy may mediate success. We find that training improves success (t(119)= 10.07, p=1.2×10^-17^), and that tDCS boosts this effect (interaction: t(119)= 2.20, p = 0.02), with no direct effect of tDCS. Stereotypy had no direct effect on success (t(119)= 0.84, p=0.74). To summarize these results, we see a direct effect of training on both stereotypy and success (Fig. 5H). We also see an effect of tDCS on stereotypy and training, however, this depends on training, as we hypothesized. Importantly, tDCS increases success (+ sign), while it reduces stereotypy (-sign).

## Discussion

We have developed a stimulation montage that allows for 10 days of stimulation without restricting the mobility of rats. This montage can serve as a standard tool for future tDCS animal experiments. We also measured the electric field in M1 and established the current intensities required to achieve the clinically comparable field magnitudes (2 V/m per 0.150 mA). This value, which is valid for this specific montage, should serve as a guideline for future translational studies.

We found that the calibrated current intensity was lower than the typical value of 250-350 µA used in previous *in vivo* experiments (Barbati et al., 2020; Cambiaghi et al., 2010; Rohan et al., 2015). In mice, 250 µA currents result in fields of 15 V/m in the hippocampus (Yu et al., 2023), and are likely stronger on the more superficial motor cortex. Thus, we argue that field intensities in these previous animal studies are much larger than in the current experiment, and likely much larger than in human tDCS experiments, where stimulation rarely exceeds 1 V/m (Huang et al., 2017; Opitz et al., 2016). Neurons in humans are larger than in rodents, which may contribute to an increase in membrane polarization due to transcranial stimulation. While somatic polarization due to electric fields has not yet been experimentally tested in human neurons, computational work using biologically realistic models of both human and rat neurons has shown that the somatic polarization lengths are quite similar, differing by only a few uV/(V/m) (Aberra et al., 2018).

Our findings demonstrate that concurrent anodal tDCS enhanced motor skill training even at small field magnitudes in rodents. We believe the key factors to achieve this were to stimulate across consecutive sessions and to apply stimulation concurrently with the training task. While we have not directly demonstrated the necessity of these factors, we note that previous studies using single-session stimulation (Cambiaghi et al., 2010) or non-concurrent (“offline”) stimulation (Barbati et al., 2020) all used higher current intensities. Previous work in animals (Sharma et al., 2022) and humans (Reis et al., 2009) support the cumulative effects found here. An *in vivo* study in mice demonstrated that multiple offline sessions of anodal tDCS can enhance context discrimination (Yu et al., 2023). In this earlier study, the current intensity was probably high and therefore the animals were lightly anesthetized during stimulation. The cumulative effect of DCS on synaptic plasticity induction was also demonstrated *in vitro* (Sharma et al., 2022). A prior human study has demonstrated that several sessions of concurrent (“online”) anodal tDCS can boost performance in an isometric pinch task (Reis et al., 2009). Finally, the importance of concurrent stimulation in the case of small field intensities had been demonstrated for *in vitro* synaptic plasticity experiments (Kronberg et al., 2017, 2020).

The specific pellet-reaching task has been explored in the context of tDCS previously. A study in mice found that offline tDCS repeated on three consecutive days improved offline learning in a test conducted 24 hours after the last tDCS session (Barbati et al., 2020). The study measured the change in performance in a single training session, which limits the ability to fully assess the cumulative effects of spaced learning. A study in rats after stroke showed an acute effect of tDCS on reaching behavior, however, it is not clear if these effects outlast the period of stimulation nor was it clear what electric field intensity was used (Ramanathan et al., 2018). A recent study in rats also demonstrated an improvement in motor recovery after stroke when using tDCS for three days after the stroke (Longo et al., 2022). To our knowledge, the present study is the first in vivo animal experiment to demonstrate an interaction between tDCS and concurrent and repeated motor learning, consistent with prior in vitro work showing an *interaction* of DCS with synaptic plasticity induction (Farahani et al., 2021; Kronberg et al., 2020).

Motor skill learning has been associated with an acute increase in neural activity during the execution of the motor task, e.g. (Olivo et al., 2022). This acute increase diminishes with practice as the tasks become more automatic (Dayan & Cohen, 2011). We have shown in vivo that the acute effect of anodal tDCS at clinically relevant field intensities is to increase firing rate (Farahani et al., 2024), and that firing rate during plasticity induction correlates with the tDCS induced boost of synaptic efficacy. (Farahani et al., 2021). What the long term effects of tDCs on firing rate are during repeated training is not yet known.

In the current study, we did not control for sensation produced by tDCS, possibly in the chest area of the cathodal return electrode. It is therefore possible that the observed effects are confounded by sensation or arousal effects, despite the lower stimulation intensities. Indeed, in right-pawed rats, the improvement in stereotypy across training sessions was reduced with tDCS. This is consistent with the sensation of stimulation disrupting the behavior of the rats. One way of addressing this potential confounding factor in future studies will be to apply cathodal tDCS in the control group as a means of controlling for sensation. Additionally, one could measure performance in the active group, in a stimulation-free period to rule out acute effects of stimulation on performance.

A second significant methodological issue comes from the control group used. In the study, the control group received electrode implants with no stimulation applied. Importantly, because the only effects observed were behavioral, this does not exclude the possibility that the stimulation was perceived by the rats, increasing their level of arousal, leading to more effective training that is unrelated to the actual mechanism of tDCS. To exclude this possibility, a control group with stimulation applied in a non-relevant area such as visual cortex should be included. Additionally, the training effects should be tested both with stimulation on and off to verify that any gains in training are maintained after the tDCS stimulation is removed.

We expected that increased stereotypy will result in better performance. However, the exploratory analysis of stereotypy and success did not confirm this. Previous studies on stereotypy and success have reported that the movement pattern of rats’ forelimbs became more similar with an increase in success rate (Lemke et al., 2021), similar to what we found here. In this earlier study, the improvement in the success rate continued after reaching a stable pattern in these forelimb movements. While Lemke et al. (Lemke et al., 2021) studied the variation in gross movement, Bova et al. (Bova et al., 2019) investigated fine motor skill comparable to our study. In that study, rats that enhanced their success rates in reaching also displayed decreased movement variability. Interestingly, even rats with low success rates showed reduced variability in their movement patterns. Together, this suggests that an increase in stereotypy is not necessarily associated with an increase in success. Rats may achieve consistency in performing a reach, even if they are unable to execute it successfully. We found that tDCS boosted gains in success rate but reduced stereotypy. But we did not find an association between stereotypy and success consistent with the results of Bova et al. One caveat to our conclusions on stereotypy is that these results are from a post hoc analysis in which we separated left and right-pawed animals, and we are not adequately powered to control for multiple comparisons of these post hoc tests. Therefore, future studies are needed to confirm these results in a new cohort of animals. Despite this caveat, the difference between left and right animals was marked. This is in contrast to previous studies which have not reported any difference between right- and left-pawed rats (Bova et al., 2019; Ellens et al., 2016; Lemke et al., 2021; Salameh et al., 2020), although there exists reports on behavioral differences with paw-preference (Ecevitoglu et al., 2020). Multiple factors may have contributed to the difference between left and right-pawed animals. For instance, left-pawed animals may be ambidextrous and therefore they used more diverse reaching strategies. It is also possible that our method to measure stereotypy may have been less accurate in left-pawed animals. Finally, the left-pawed animals performed relatively poorly compared to previous studies [41,52], which may have limited the effects of tDCS. However, we hesitate to make strong claims given that these were exploratory analyses without multiple comparison corrections.

Two final caveats on the learning process are as follows: On average, the success rate in our experiment was close to the success rate reported in previous studies where the rats were fed automatically (Bova et al., 2019; Ellens et al., 2016). We also restricted the duration of each tDCS to 20 minutes, aligning with the customary application of tDCS in human experiments. We aimed to maintain consistency in the duration of both training and stimulation for comparison. This has limited the number of attempts during a session and may have impacted the learning progress. Additionally, the rat had to learn to move back and forth to fetch a new pellet from the pedestal on the front panel. As a result, the learning process involves two components: reaching, which requires dexterous motor skill learning, and a cognitive component, whereby the animal has to learn that the pedestal only reloads when going to the back of the chamber. It is not clear which of these components were affected by tDCS.

In humans, MEPs are evoked by single-pulse TMS. Previous studies on the effect of tDCS on these TMS-MEP have produced inconsistent results. While some studies have demonstrated that tDCS can modulate TMS-MEPs (Nitsche & Paulus, 2000; Rosenkranz et al., 2007; Ziemann et al., 2004), others have failed to find any significant effects (Horvath et al., 2015; Jonker et al., 2021). In our study, we found that MEPs are stronger in the trained paw in the control group, but we did not see the expected increase in MEP with tDCS. This suggests that the mechanisms affecting MEPs are different from those affecting skill learning, or that our study was underpowered to detect smaller tDCS effects on MEPs.

MEP amplitude is a marker of corticospinal excitability (Pellicciari et al., 2013), whereas learning dexterous control may rely on system-wide synaptic connections and have little to do with simple increased excitability, which is affected instead by strength training for instance (Adkins et al., 2006). In this context, it is worth noting that we have found no reports to date of increased MEP amplitude with motor learning in animals or humans. What has been reported is an expansion of the motor cortex map with the same reaching task and using a similar MEP protocol (Brus-Ramer et al., 2009). In humans, motor map expansion (measured with TMS-MEP) has been demonstrated with a task more akin to repetitive strength training rather than fine motor skill (Wang et al., 2021). With our montage there may be significant current flow along the corticospinal tract, which would not be lateralized. If electric fields affect excitability in the spinal track, then this might explain the lack of specificity to the trained paw. Future studies could shed light on whether this nonspecific effect is also observed in behavior, such as whether reaching performance is also improved in the untrained paw. A recent human study has shown unspecific effects of tDCS on motor skill learning, improving both online and offline learning as well as the stimulated and unstimulated hemisphere (Hsu et al., 2023).

In conclusion, we presented a stimulation montage that allows repeated learning experiments across multiple days in freely behaving rats. The results are consistent with the basic hypothesis that tDCS acts as a modulator of learning, rather than improving motor skill on its own. We observed this for reaching success, in a planned analysis, and for stereotypy in an exploratory analysis. Contrary to our expectation, however, we did not detect consistent MEP effects of tDCS. These suggest that these outcomes may have a different physiological substrate than the behavioral effects reported here. Or more simply, what we were not adequately powered to detect those effects. We hypothesized that effects are mediated by a boost of synaptic plasticity, but we cannot rule out effects of sensation or arousal, among others. Ultimately, however, we have shown a robust effect in a model system that now allows systematic analysis of the mechanism by which tDCS affects motor learning.

## Acknowledgments

This work has been supported by NIH through NIH R01NS130484 and R01NS095123. We express our gratitude to Neela Zareen and Hisham Sharif for initial guidance on rodent surgeries, and Niranjan Khadka and Marom Bikson for their help on estimating field intensities based on current-flow modeling.

## Extended Data

**Figure 4-1: Motor evoked potential (MEP) measurements.** Left: Raw MEP signals (stimulation artifacts truncated) for 20 trials (color cycle) 10 ms after the last pulse of stimulation. Middle: rectified MEP. Right: median across repeats, here for the 140% condition stimulation. A/B/C are for three different animals.

**Figure 5-1: Summary of statistical analysis including paw-preference as a factor.** Links indicate significant regression parameters (p<0.05) in two separate linear mixed effect models, one for the number of successes and another for stereotypy as dependent variables. tDCS, training day and paw were the dependent variables (fixed effects), while animal was a random effect variable. Missing arrows indicate non-significant regression parameters (p>0.05). Not all possible 3-way interactions were tested. We caution that these post hoc analyses are not corrected for multiple comparisons and are not adequately powered. They should be considered exploratory and would need to be confirmed in new planned experiments.

**Figure 5-2: Reaching trajectories for four digits. A**: The trajectory data for a single grab video, covering the entire 400 ms duration. **B:** The template trajectory, derived from the average grab across three trials, specifically at the time of the grab (150 ms). **C:** The cross-correlation analyses were conducted for each digit compared to the template grab, with an average computed across all digits. The peak correlation is utilized to predict the location of the grab (30ms). **D:** The detected grab location shown in the original trial (30 ms - 180 ms). **E:** The predicted grab locations separately within the time window of 30 ms to 180 ms.

