## Supplementary figures and images for "Repeated tDCS at clinically-relevant field intensity can boost concurrent motor learning in rats"

### Supplementary Figure 1

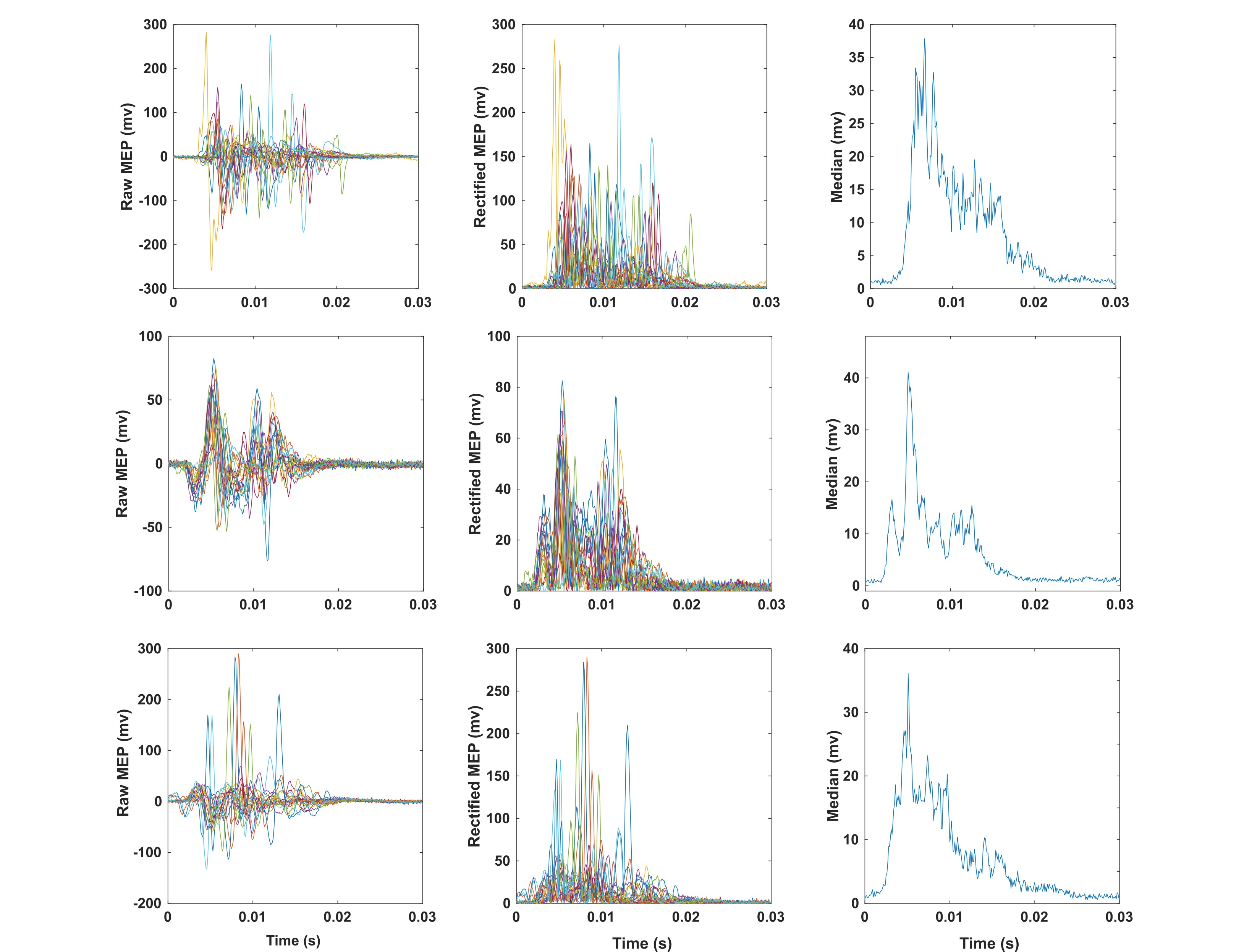

### Supplementary Figure 2

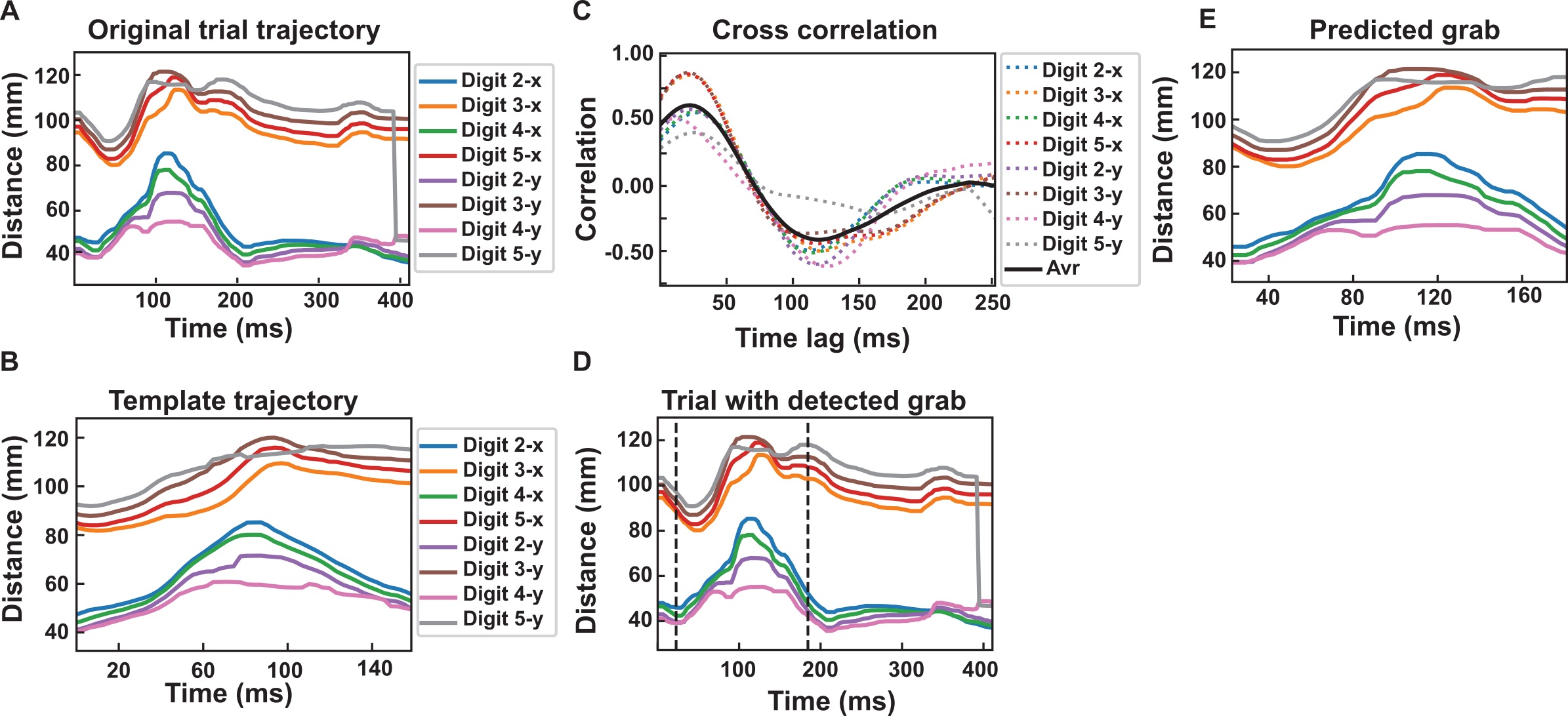

### Supplementary Figure 3

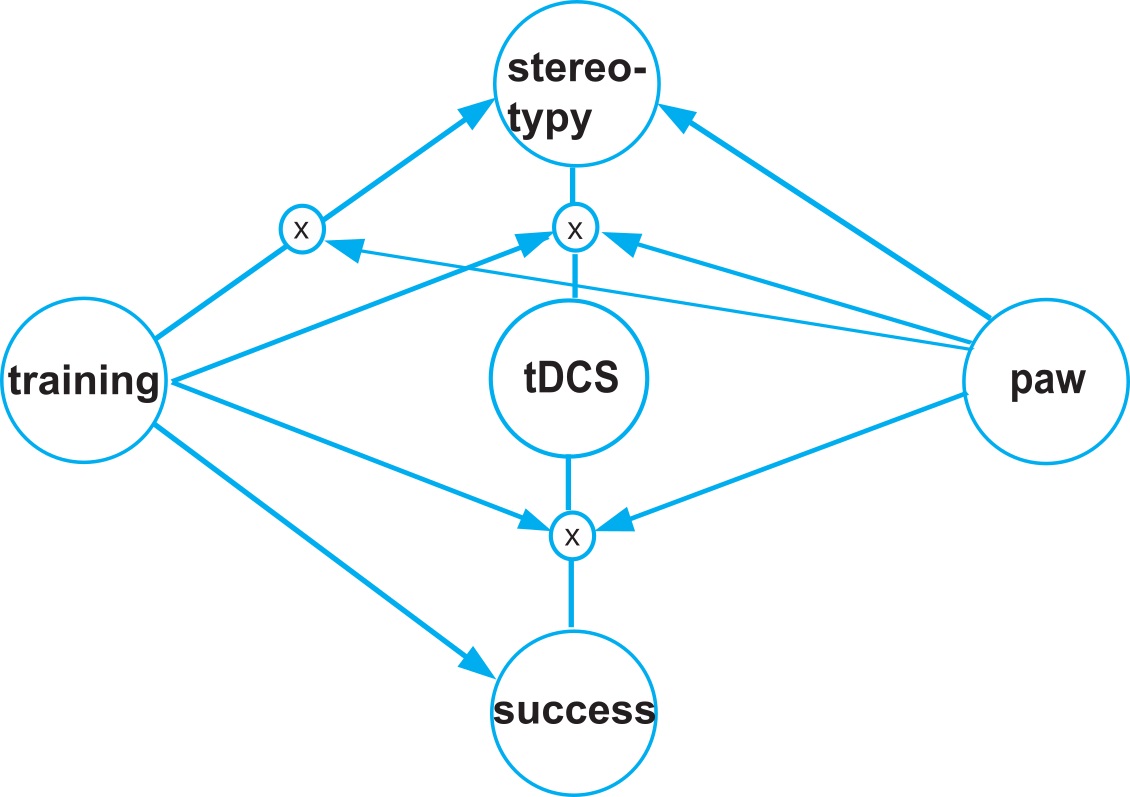
